# The effect of norepinephrine on ovarian dysfunction by mediating ferroptosis in mice model

**DOI:** 10.1101/2024.07.30.605932

**Authors:** Hanqing Hong, Chengqi Xiao, lichun Weng, Qian Wang, Dongmei Lai

## Abstract

Studies shows that stress is associated with ovarian dysfunction. Norepinephrine (NE), a classic stress hormone in the stress response, is less recognized for its role in ovarian function. A NE-treated mouse model is induced by intraperitoneal injection of NE for 4 weeks. Compared with the normal control, we find that NE-treated mice show disturbances in the estrous cycle, decreased levels of anti-Mullerian hormone (AMH) and estradiol (E2), and increased levels of follicle-stimulating hormone (FSH). Additionally, the number of primordial follicles, primary follicles, secondary follicles, and antral follicles decreased, while the number of atretic follicles increased in NE-treated mice, indicating NE-induced ovarian dysfunction. RNA sequencing further reveals that genes associated with ferroptosis are significantly enriched in NE-treated ovarian tissues. Concurrently, the content of reactive oxygen species (ROS), ferrous ion, and malondialdehyde (MDA) increased, while the expression level of glutathione peroxidase 4 (GPX4) decreased.

To elucidate the mechanism of NE-induced ferroptosis in ovaries and the potential reversal by Coenzyme Q10 (CoQ10), an antioxidant, we conduct both in vitro and in vivo experiments. In vitro, we observe that the granulosa cell line KGN, when treated with NE, shows decreased cell viability, reduced expression of GPX4, elevated ferrous ion and ROS content, and increased MDA levels. However, these NE-induced changes are rescued by the addition of CoQ10. In the mouse model, we find that NE-treated mice supplemented with CoQ10 increased GPX4 levels and decreased the contents of iron, ROS, and MDA compared with the NE group. Moreover, the differential expression of genes associated with ferroptosis induced by NE is ameliorated by CoQ10 in NE-treated mice. Additionally, CoQ10 improved ovarian function, as evidenced by increased ovarian weight, more regular estrous cycles, and an increase in follicles at various stages of growth in NE-treated mice. In conclusion, NE induces ovarian dysfunction by triggering ferroptosis in ovarian tissues, and CoQ10 represents a promising approach for protecting reproductive function by inhibiting ferroptosis.

## Introduction

Premature ovarian insufficiency (POI) refers to the decline in ovarian function in women before the age of 40 years, characterized by elevated gonadotropin levels and fluctuating low estrogen levels. The global incidence is approximately 3.5% and is on the rise[1-3]. This condition leads not only to infertility but also to severe health consequences, including hot flashes, depression, headaches, cognitive decline, vaginal atrophy, cardiac risks, and osteoporosis[1].

The etiology of POI is complex, with an increasing number of studies suggesting that stress plays a crucial role in human reproductive health. Clinical studies have demonstrated that stress can lead to abnormal reproductive function[4, 5]. For instance, a study evaluating 7352 women attending infertility clinics found that 56% exhibited significant depressive symptoms and 76% showed significant anxiety symptoms[6]. The impact of stress on reproductive function is multifaceted, including inhibition of ovulation, reduction of fertilization rate, and hindrance of fertilized egg implantation[7]. Recent research from our group revealed that POI patients experienced a higher incidence of adverse life events related to work stress, family stress, and sleep problems before diagnosis, suggesting that these patients might have been exposed to long-term chronic stress prior to illness onset[8]. Stress can activate the hypothalamic-pituitary-adrenal axis, leading to prolonged abnormal endocrine hormone levels in women, ultimately resulting in menstrual disorders, infertility, and other reproductive dysfunctions[9]. In animal experiments, chronic stress has been shown to disrupt the estrous cycle, alter hormone levels, and decrease follicle numbers in mice[10, 11]. Stress can also induce cytoplasmic fragmentation, apoptosis, spindle disorders, and oxidative stress, accelerating the aging of mouse oocytes and increasing the risk of early pregnancy failure[12].

Norepinephrine, a key catecholamine neurotransmitter involved in stress signaling, has been implicated in various psychological disorders resulting from stress, including cardiovascular disease, affective disorders, post-traumatic stress disorder, and cancer[13]. Evidence has shown that NE is involved in the regulation of ovarian functions. NE is linked to ROS-regulated events in ovarian physiology, including ovulation[14]. Increased NE content has been found in the follicular fluid of patients with polycystic ovary syndrome[15]. Wang et al. reported that glutamine and norepinephrine in follicular fluid synergistically enhance the antioxidant capacity of human granulosa cells and may predict the outcome of in vitro fertilization and embryo transfer[16]. Although NE plays an important role in the physiology and pathology of the ovary, the effects of excessive NE induced by chronic stress on ovarian function are not well understood. This study aims to establish a NE-treated mouse model, explore the mechanism of excessive NE on ovarian dysfunction, and preliminarily screen potential protective agents against ovarian injury induced by chronic stress.

Ferroptosis is an iron-dependent, non-apoptotic mode of cell death characterized by lipid peroxide accumulation. In 2012, Professor Brent. R. Stockwell of Columbia University proposed the concept of ferroptosis for the first time[17]. When there is iron metabolism disorder in cells, iron accumulates in cells, ROS level increases, lipid peroxidation occurs in cells, lipid peroxidation products cannot be effectively removed, accumulate in cells, mediate cell membrane damage, and ferroptosis occurs in cells[18, 19]. The occurrence of ferroptosis is closely related to ovarian dysfunction. Ovary bioinformatics analysis in mice, the results showed that iron death marker protein TFRC, NCOA4 and SLC3A2 in aging expression is significantly higher than young group[20]. BNC1 deficiency leads to follicle activation and excessive follicle atresia while triggering oocyte ferroptosis through NF2-YAP pathway, and mice exhibit ovarian dysfunction symptoms[21]. Studies show that overexpression of miR-93-5p can negatively regulate the NF-κB signaling pathway to promote granular cell death of iron, and knocked down the miR-93-5p can save granulosa cell function[22]. After dehydroepiandrosterone treatment, the expression of transferrin receptor 1 and iron content in mouse ovaries increased, which mediated the release of reactive oxygen species, activated mitophagy, induced lipid peroxidation, and promoted ferroptosis of granulosa cells[23]. The expression levels of antioxidant genes, steroid biosynthesis gene, and ferroptosis marker (GPX4) were down-regulated in ovarian tissues and KGN cells of cigarette smoke-treated mice, suggesting that cigarette smoke may cause ferroptosis in ovarian granulosa cells, leading to reduced fertility in mice[24]. Therefore, ferroptosis plays an important role in the development of ovarian dysfunction. An in-depth study of the relationship between ferroptosis and ovarian function provides new ideas for the prevention and treatment of ovarian dysfunction.

## Materials and methods

### Animals

Eight-week-old C57BL/6 female mice were obtained from Shanghai Jiao Tong University School of Medicine. Mice were maintained in the Department of Animal Experimentation, School of Medicine, Shanghai Jiao Tong University. Adaptation in a controlled laboratory (25 ° C, 55±5% relative humidity, 12 h light/dark cycle) for 1 week. Animals were maintained in mouse cages and had free access to food and water throughout the study period. This study was a controlled experimental study on experimental animals. Ethical approval for this study was obtained from the Department of Animal Science, Shanghai Jiao Tong University School of Medicine (IPMCH GKLW 2022-61). All animal experiments were performed in strict accordance with the guidelines for the management and use of animal experiments.

### Experimental design

In animal experiments, 8-week-old female C57BL/6 mice were randomly divided into four groups: Ctrl group (control, saline, intraperitoneal injection), NE group (intraperitoneal injection, 1mg/kg/ day), NE (intraperitoneal injection, 1mg/kg/ day) + CoQ10 (100mg/kg/day, feeding) group and CoQ10 group (100mg/kg/day, feeding,). The mice were treated for 4 weeks. In cell experiments, four groups were divided. Ctrl group, NE (1000μM) group, NE (1000μM) + CoQ10 (1μM) group, and CoQ10 (1μM) group. CoQ10 and NE were purchased from MCE company (Shanghai, China).

### RNA-sequencing (RNA-seq) experiments and analysis

Mouse ovaries were divided into four groups for transcriptome sequencing analysis. The four groups were the control group, NE group, NE + CoQ10 group, and CoQ10 group. Transcriptome sequencing and analysis were performed by Shanghai Ou Yi Biotechnology Company (Shanghai, China). DESeq2 software was used to analyze differentially expressed genes, which were consistent with q value< 0.05 and foldchange> 1.5. Genes with a threshold of 1.5 were defined as differentially expressed genes (DEGs). Kyoto Encyclopedia of Genes and Genomes (KEGG) Pathway enrichment analysis of differentially expressed genes was performed based on a hypergeometric distribution algorithm to screen significant enriched functional items. R (v 3.2.0) was used to draw an enrichment analysis map for significant enrichment functional items. Gene set enrichment analysis was performed using GSEA software. The predefined gene sets were used to rank the genes according to their degree of differential expression in the two types of samples, and then to test whether the prespecified gene sets were enriched at the top or bottom of the ranking table.

### Hematoxylin and eosin staining (HE) and follicle count

Mouse ovaries were fixed in 4% paraformaldehyde for more than 24 hours. Ovaries were dehydrated, embedded in paraffin, serially cut into 5 μm sections, and placed on slides. dewaxing solution (1) for 20 min, dewaxing solution (2) for 20 min, absolute ethanol (1) for 5 min, absolute ethanol (2) for 5 min, 75% alcohol for 5 min, and tap water for 1 min. It was placed in a hematoxylin staining solution for 1-2 min and rinsed with running water. In return blue solution for 1-2 min and rinse with running water. The cells were stained in an eosin staining solution for 3 minutes. The slices were rinsed with running water, followed by absolute ethanol (1) for 5 min, absolute ethanol (2) for 5 min, deparaffinized transparent solution (1) for 5 min, deparaffinized transparent solution (2) for 5 min, and sealed with neutral gum. Image acquisition and analysis were performed after microscopic observation (LEICA DMi8). Follicle counts for each section were done by two observers who were unaware of the source of the sample. The number of primordial, primary, secondary, antral follicles, and atretic follicles was calculated. Primordial follicles are oocytes surrounded by a flat layer of granulosa cells. A layer of cuboid granulosa cells surrounds the oocytes of the primary follicle. Secondary follicles consisted of an oocyte and surrounded by more than one layer of granulosa cells, with no visible antrum. Antral follicles are follicles in which the antral cavity can be seen. All reagents were purchased from Servicebio Company (Wuhan. China).

### Enzyme-linked immunosorbent assays (ELISA)

After the mice were anesthetized, eyeball blood was collected, and serum was obtained after standing and centrifugation. The levels of AMH, FSH, and E2 in serum were detected by ELISA kit. The absorbance value was measured at 450 nm, and the serum concentrations of AMH, FSH, and E2 were calculated. Statistical analysis was performed using GraphPad Prism 8 software. ELISA kits were purchased from CUSABIO Company (Wuhan, China), and all procedures were performed in strict accordance with the instructions.

### Estrous cycle examination

The vaginal smear test was performed from 9 a.m. to 10 a.m. each day. Estrous cycles were determined by Giemsa staining. 10 μL saline was injected into the vagina. The aspiration was performed three times. Saline was applied to the slide. After drying, staining was performed using Giemsa staining solution. The estrous cycle was observed and recorded under a light microscope. The proportion of nucleated, keratinized epithelial cells and leukocytes determines the type of estrous cycle. Proestrus: abundant nucleated epithelial cells, few keratinized cells. Estrus: Numerous keratinized epithelial cells, few nucleated. metestrus: mainly non-nucleated keratinocytes and neutrophils, with a higher proportion of neutrophils. Diestrus: fewer cells, neutrophils predominate. Disturbance of the estrous cycle is a prominent feature of ovarian dysfunction. Giemsa staining kit was purchased from Beyotime (Shanghai, China).

### Immunohistochemistry

The ovarian tissue was cut into 5 μm. Deparaffinized and rehydrated in deparaffinized solution and ethanol of different concentrations. Antigen repair solution (Servicebio, Wuhan, China) was used for antigen repair. Endogenous peroxidase was blocked by washing with TBS three times for 1 minute each time. The cells were blocked in a rapid blocking solution (Beyotime, Shanghai, China) for 10min. Primary antibody was added and incubated at room temperature for 1h. The cells were washed three times with TBS (Beyotime, Shanghai, China) for 1 minute each time. Secondary antibodies were added and incubated at room temperature for 30 minutes, then washed thrice with TBS for 1 minute each time. DAB was used for color development (Servicebio, Wuhan, China). Nuclei were stained in hematoxylin solution. Neutral gum was used for sealing. Photographs were taken and analyzed. Image J software and GraphPad Prism 8 software were used to perform staining intensity analysis and statistical analysis. Antibodies used were as follows: GPX4 (1:2000, protein tech); Goat Anti-Mouse secondary antibody (Ready-to-use, proteintech, PR30012).

### Cell culture

Granulosa cell line KGN cells were cultured using DMEM/F12 supplemented with 10% fetal bovine serum and 100 IU/mL penicillin, 100 μg/mL streptomycin as a complete medium. Cells were cultured in a constant temperature incubator (5% CO2) at 37°C. The ovarian granulosa cell tumor cell line (KGN) was purchased from Shanghai Fuheng Biotechnology Company (FH1125). STR identification was performed on KGN cells.

### Cell viability assay

Cell viability was measured by CCK8. Cells were seeded in 96-well plates (100 μL/well). After the cells were adherent, drugs were added and the cells were incubated for 48 hours. 10 μL of CCK8 solution were added to each well. The 96-well plates were placed in the incubator and incubated for 2 hours. The absorbance at 450 nm was measured using a microplate reader. GraphPad Prism 8 software was used for statistical analysis of the data. The drugs used were as follows: NE (1000 μM) and CoQ10 (1μM). The above reagents were purchased from the MCE company (Shanghai, China).

### Western Blotting

Preparation containing protease inhibitors and phosphatase inhibitory RIPA pyrolysis liquid, placed in 4 □ for later use. The cell culture medium was removed, washed twice with PBS, and digested with 0.25% trypsin for 2-3 minutes. The digestion was terminated using the medium in DMEM/F12 containing 10% FBS. The bottom of the cell culture dish was gently blown to suspend the cells, and the cell suspension was transferred to a centrifuge tube, placed in a centrifuge, and centrifuged at 1500 rpm for 5 minutes. The supernatant was discarded and 200 µL of lysate containing RIPA with protease inhibitors and phosphatase inhibition was added. They were placed on ice and allowed to lyse for 30 min, vortexed every 10 min for 1 min each time. After completion of lysis, the cells were placed in a centrifuge and centrifuged at 1500 rpm for 5 min. The supernatant was transferred to a new centrifuge tube for protein concentration determination. The BCA method is used to detect protein concentration. Fix the polyacrylamide gel electrophoresis tank, filled with electrophoresis between liquid to the gel plate, pull out the tooth comb, the protein marker, and samples under test to the corresponding lane, for constant voltage electrophoresis, 110 v electrophoresis for 50 minutes, stay bromophenol blue close to stop at the bottom of the gel electrophoresis. The gel plate was removed, and after rinsing with water, the glass plate was pricked to remove the gel and placed in the transfer solution. The PVDF membrane was activated by immersing it in methanol for 10 seconds. The film clip was made in the following order: black film clip - sponge - filter paper - gel -PVDF membrane - filter paper - sponge - white film clip. Transfer the film clip to the film tank, add the film transfer solution, and put in ice cubes. Constant flow was performed and the membrane was turned at 200 mA for 90 minutes. The PVDF membrane was immersed in a blocking solution and blocked for 15 min at room temperature. Primary antibodies were incubated overnight at 4 ° C. The primary antibody was recovered and the PDVF membrane was washed three times for 5 min each using TBST on a shaker. The cells were incubated with horseradish peroxidase-labeled secondary antibodies for 1 h at room temperature. Secondary antibodies were recovered, and PDVF membranes were washed three times for 5 min each using TBST on a shaker. The ECL chemiluminescence kit was used for visualization. Image J software was used to quantify protein bands. The relative protein expression was calculated using an internal control for standardization. GraphPad Prism 8 (GraphPad Software, San Diego, CA, USA) software was used for statistical analysis. The antibody used was as follows: GPX4 (1:5000, Proteintech,67763-1-Ig); Goat Anti-Mouse secondary antibody (1:4000, Abcam, ab6789).

### Measurement of ovarian tissue ferrous ion content

After the mice were anesthetized, ovarian tissues were collected. Ovarian tissue and extracts were mixed and homogenized in an ice bath. Placed in a centrifuge, the temperature was set to 4 ° C, centrifuged at 4000 g for 10 min, and the supernatant was removed. The microplate reader was preheated for 30 min and the wavelength was adjusted to 520 nm. After sample addition, the samples to be tested were thoroughly shaken and mixed, centrifuged at 10000 rpm at room temperature for 10 min, and 200 μL of the upper inorganic phase was carefully sucked, transferred to a 96-well plate, and the absorbance was measured at 520 nm. Kits were purchased from Beijing Solarbio Science & Technology Company (BC5415).

### Measurement of cellular ferrous ion content

The cells were added to the extract and mixed (0.5ml of the extract was added per 5 million cells). The cells were broken by ultrasound (power 200 W, 3 seconds, interval 7 seconds, repeated 30 times) in an ice bath, centrifuged at 8000 g for 10min at 4□, and the supernatant was taken for testing. The microplate reader was preheated for 30 min and the wavelength was adjusted to 510 nm. The samples to be tested were added to a 96-well plate and thoroughly mixed, left at 25 □ for 10min for color development, and the absorbance value at 510 nm was measured in a 96-well plate. Kits were purchased from Beijing Solarbio Science & Technology Company (BC5415).

### Tissue reactive oxygen species (ROS) content analysis

Reagent preparation: The O11ROS probe was diluted 10-fold using pure water. Mix well and set aside. Sample processing: The obtained fresh ovarian tissue was cleaned with PBS. Ovaries were placed in homogenization buffer A at a mass (mg) to volume (μL) ratio of 1:10. Homogenate as fully and rapidly as possible. They were placed in a centrifuge at 100xg for 5 min at 4 ° C, and the supernatant was removed for use. Protein quantification: Tissue homogenates were diluted 30-fold in PBS before protein quantification. Sample detection: Homogenate supernatant and O11ROS probe (19:1 volume ratio) were added to a 96-well plate and blown with a pipette to allow thorough mixing. The plates were incubated at 37 ° C in the dark for 25 min. The fluorescence intensity was detected at an excitation wavelength of 488 nm and an emission wavelength of 530 nm in a microplate reader. The reactive oxygen species detection kit (BB-470522) was purchased from Bebo Biological Company.

### Cell reactive oxygen species (ROS) content analysis

KGN cells were seeded in 6-well plates. When the cells grew to 70%, the control group was replaced with a complete medium, and the treatment group was replaced with a medium containing NE (1000 μM). The cells were incubated for 48 hours. DCFH-DA was diluted in a serum-free medium at a volume ratio of 1:1000 to achieve a final concentration of 10 μL. The cell culture medium was removed, and the diluted DCFH-DA diluent was added after the cells were collected. The cells were suspended in the diluted DCFH-DA diluent and incubated in a cell incubator at 37 □ for 20 minutes. The mixture was reversed at intervals of 3-5 min to allow adequate contact between the probe and the cell. Cells were washed three times with serum-free cell culture medium to adequately remove DCFH-DA that did not enter the cells. Fluorescence intensity was measured using a microplate reader with an excitation wavelength of 488nm and an emission wavelength of 525 nm. The ROS detection kit was purchased from Beyotime Biotechnology Company (Shanghai, China).

### Measurement of MDA content

Cell sample preparation: cells were collected into a centrifuge tube, and the supernatant was discarded after centrifugation. 1mL of the extract was added per 5 million cells, and the cells were broken by ultrasound (power of 200W, 3s, interval of 10s, repeated 30 times). After centrifugation at 8000g for 10 min at 4°C, the supernatant was removed and placed on ice until measured. Preparation of tissue samples: 1mL of extract was added to every 0.1g of tissue. Ice bath homogenate. After centrifugation at 8000g for 10min at 4 °C, the supernatant was removed and placed on ice until measured. The microplate reader was preheated for more than 30 minutes, and distilled water was zeroed. The MDA working solution, distilled water, and samples were mixed. After the mixture was kept in a water bath at 100 ° C for 60 minutes (the cover was tight to prevent water loss), it was cooled in an ice bath at 10000 g at room temperature and centrifuged for 10 minutes.200 μL of the supernatant was aspirated in a 96-well plate, and the absorbance of each sample was measured at 532 nm and 600 nm. The MDA detection kit was purchased from Beijing Solarbio Science &Technology Company.

### Statistical analysis

Data was shown as mean ± standard error. Statistical analysis was performed via GraphPad Prism 8 (GraphPad Software, San Diego, CA, USA). Student’s t-test was utilized to compare the mean between two groups, and one-way analysis of variance (ANOVA) was used to compare three or more groups. If the data did not conform to the normal distribution, the nonparametric test was used for analysis. A difference at p<0.05 was considered statistically significant.

## Results

### 1. NE causes ovarian dysfunction in mice

To explore the effect of NE on ovarian function, C57BL/6J mice were treated with NE via intraperitoneal injection for 4 weeks (Fig. 1A). Compared to control mice, NE-treated mice exhibited significantly lower ovarian weight (Fig. 1B). Additionally, NE-treated mice showed disrupted estrous cycles, characterized by shortened estrus and prolonged diestrus (Fig. 1C, D). Ovarian morphology was examined through HE staining (Fig. 1E), revealing a decrease in the number of primordial follicles, primary follicles, secondary follicles, and antral follicles, along with an increase in atretic follicles in NE-treated mice ovaries (Fig. 1F). Furthermore, levels of AMH and E2 were significantly lower, while FSH levels were significantly higher in the serum of NE-treated mice compared to the control group (Fig. 1G, H, I). These findings indicate that NE caused ovarian dysfunction in mice.

**Figure 1.**
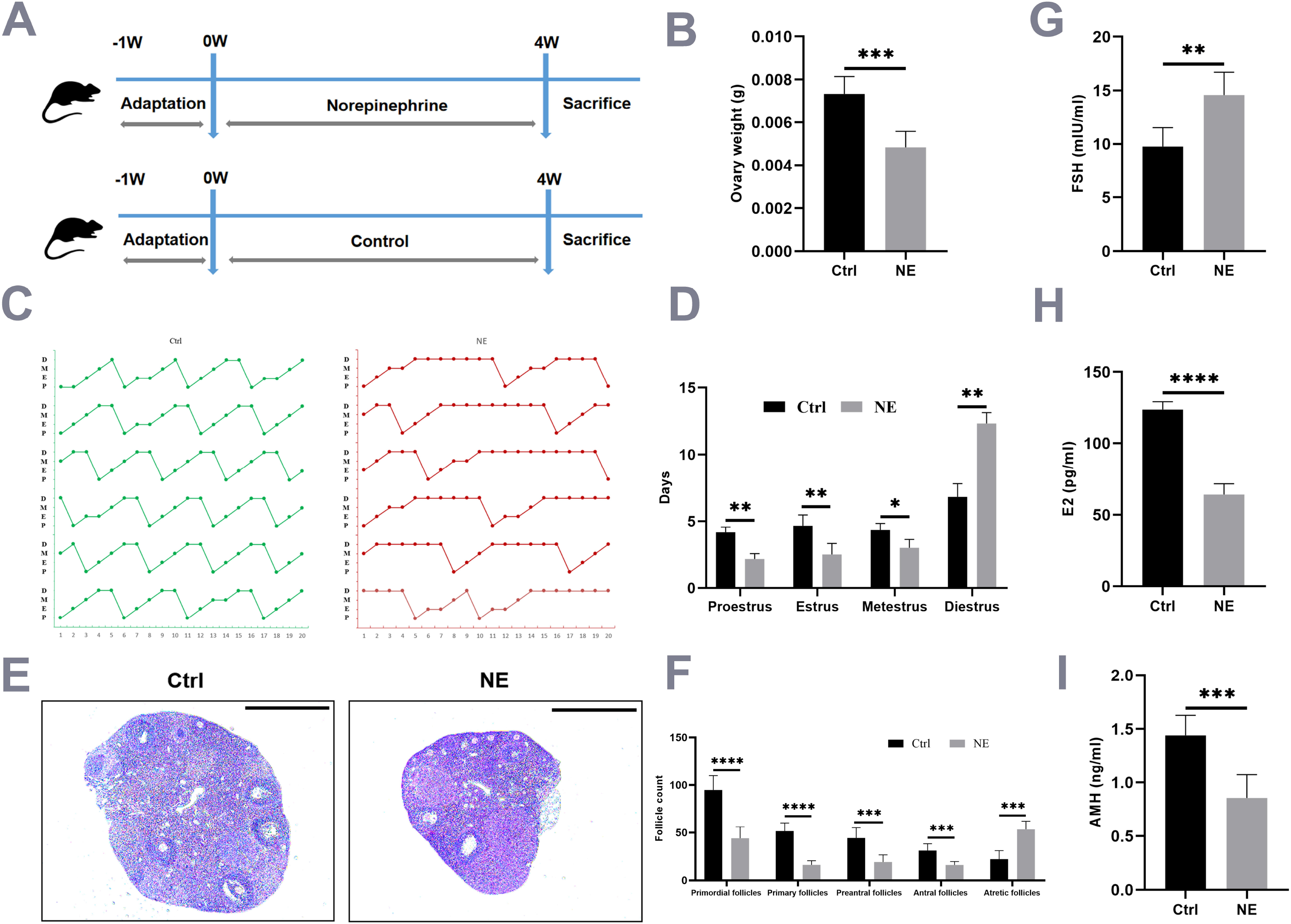
NE causes ovarian dysfunction. **A**. Experimental design of the mouse model. Female C57BL/6 mice at 8 weeks of age were randomly divided into two groups. One group of mice was injected intraperitoneally by NE (1mg/kg/d) for 4 weeks. The control group mice received intraperitoneal injections of saline for 4 weeks (n = 6). **B**. Compared with the control group, the ovarian weight of mice in the NE group was decreased (n = 6). **C, D**. The vaginal smear experiment was conducted to examine the estrous cycle. Compared with the control group, the estrous cycle in the NE group was disrupted, primarily manifested as prolonged diestrus stages (n = 6). **E**. HE staining of ovaries from control and NE mice (n = 6, bar = 500 μm). **F**. Compared with the control group, the NE group had a decrease in the numbers of primordial follicles, primary follicles, secondary follicles, and antral follicles, while the number of atretic follicles increased by follicle counting (n = 6). **G.H.I**. The hormone levels (FSH, E2, and AMH) in mouse serum were performed by ELISA. Compared with the control group, The levels of AMH and E2 were decreased, while the level of FSH was increased in the NE group (n = 6).The data were presented as mean ± S.E.M (*pC<C0.05,**p < 0.01,***p < 0.001,****p < 0.0001).Ctrl, control; NE, norepinephrine; AMH, anti-Mullerian hormone; FSH, follicle stimulating hormone; E2, estradiol.

### 2. NE induces ferroptosis in mice ovaries

To explore the mechanism of ovarian dysfunction induced by NE, we conducted transcriptome sequencing on control and NE-treated ovaries. The RNA sequencing (RNA-Seq) results revealed a significant enrichment of genes associated with ferroptosis in ovarian tissues of the NE group compared to the control group (Fig. 2A). Iron accumulation is a known key factor leading to ferroptosis[25]. Therefore, we examined the amount of iron in ovarian tissue. The Ferrous ion (Fe^2+^) assay results showed a significant elevation in the NE group compared to the control group (Fig. 2B). Additionally, the detection of ROS showed that the ROS content in the ovaries of mice in the NE group was significantly higher than that in the control group (Fig.2C). Lipid peroxidation is another key factor inducing ferroptosis. Malondialdehyde (MDA) is a highly reactive aldehyde and one of the decomposition products of lipid peroxidation, often used as a marker of lipid peroxidation[26]. Our results showed that the content of MDA in the ovaries of the NE-treated group was significantly higher compared to the control group (Fig. 2D). Glutathione peroxidase 4 (GPX4) is an important antioxidant enzyme that removes intracellular lipid hydroperoxides and protects the cell membrane from oxidative damage. Inhibition of GPX4 activity or down-regulation of its expression can lead to a loss of effective defense against lipid peroxidation, promoting ferroptosis[27]. We measured the expression level of GPX4 in ovarian tissues, which showed a significant reduction in the NE-treated group compared to the control group (Fig. 2E, F). Overall, these results suggest that NE causes ferrous ion accumulation and lipid peroxidation, ultimately leading to ferroptosis in mouse ovarian tissues.

**Figure 2.**
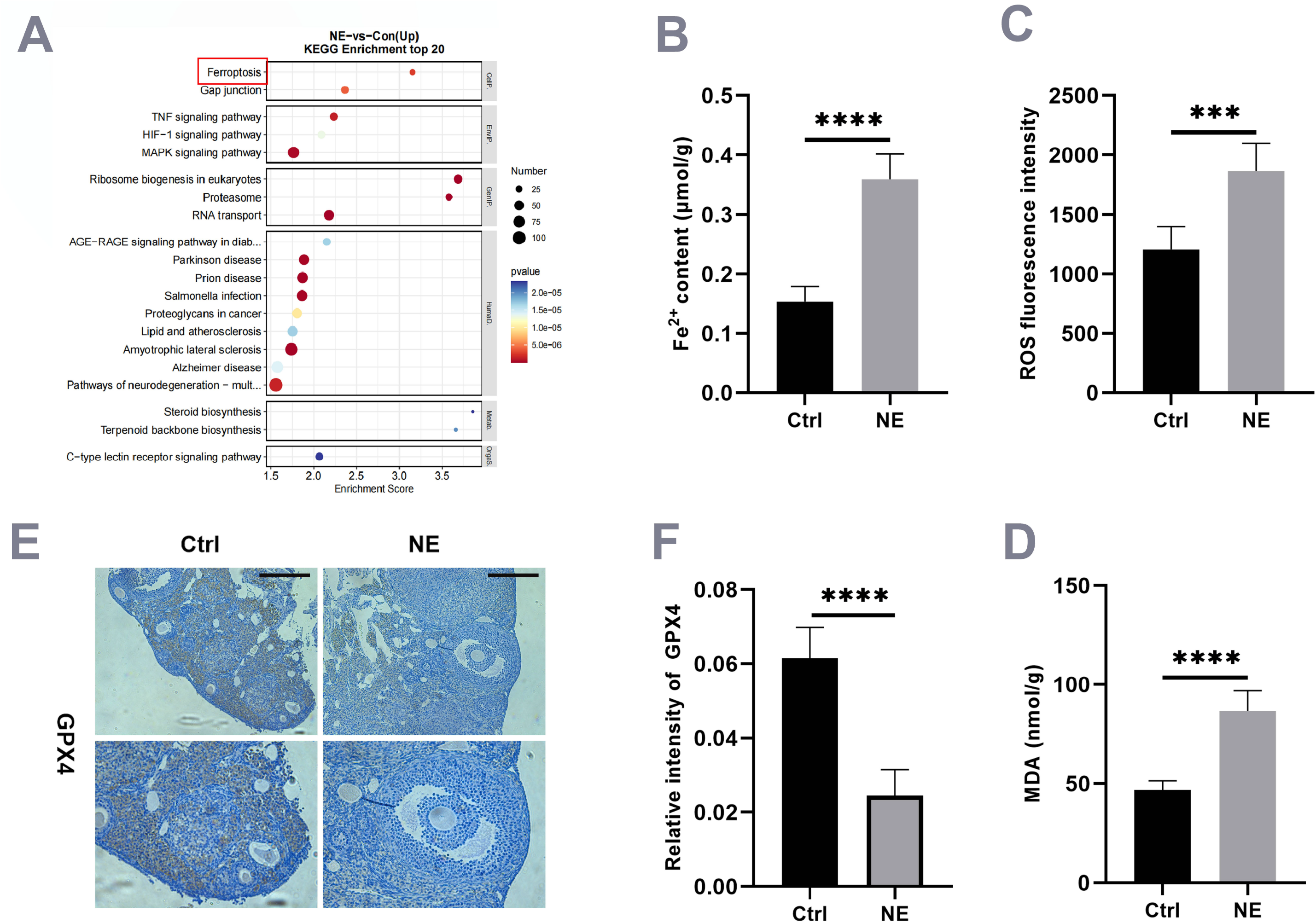
NE induces ferroptosis in mouse ovaries. **A**. KEGG analysis showed that ferroptosis-related genes were significantly enriched in the ovaries of the NE group (n = 3). **B**. Fe^2+^ assay results showed that the NE group was significantly elevated compared with the control group (n = 6). **C**. ROS assay results showed that ROS content was increased in the NE group compared with the control group (n = 6). **D**.MDA was significantly increased in the NE group compared with the control group (n = 6). **E.F**. The IHC experiment showed that the expression level of GPX4 in the ovaries of the NE group was decreased compared with the control group (n = 6, bar=200 μm).The data were presented as mean ± S.E.M (*p□<□0.05,**p < 0.01,***p < 0.001,****p < 0.0001).Ctrl, control; NE, norepinephrine; KEGG, Kyoto Encyclopedia of Genes and Genomes; MDA, malondialdehyde; GPX4, glutathione peroxidase 4; IHC, Immunohistochemistry.

### 3. CoQ10 rescues ferroptosis induced by NE in vitro

Coenzyme Q10 is a lipophilic free radical trapping antioxidant that detoxifies lipid peroxyl radicals and has been found to inhibit ferroptosis[28]. To further explore the mechanism of ferroptosis induced by NE and the potential rescue by CoQ10, we treated the ovarian granulosa cell line KGN with NE and CoQ10. The results showed that treatment with NE (1000 μM) for 48 hours reduced cell viability, as assessed by CCK8 assay. However, cell viability was significantly rescued in the NE (1000 μM) + CoQ10 (1μM) group compared to the NE (1000 μM) treated group (Fig. 3A). We also examined the effect of NE on ferrous ion (Fe^2+^) content in the cells. The results showed a significant increase in Fe^2+^ content in the NE (1000 μM) group compared to the control group, indicating that NE leads to the accumulation of ferrous ions (Fe^2+^) in the cells. However, compared to the NE group, ferrous ion (Fe^2+^) content was significantly decreased in the NE (1000 μM) + CoQ10 (1 μM) group (Fig. 3B). Furthermore, the results of ROS detection showed a significant increase in ROS content after NE (1000 μM) treatment of KGN cells, while treatment with NE (1000 μM) + CoQ10 (1μM) reduced ROS content (Fig. 3C). Additionally, we measured the level of MDA to explore whether NE could cause lipid peroxidation in cells. The results indicated that NE (1000 μM) treatment for 48 hours significantly increased the level of MDA in KGN cells. However, compared to the NE group, MDA content was significantly decreased in the NE (1000 μM) + CoQ10 (1 μM) group (Fig. 3D). Lastly, we examined the content of GPX4 in KGN cells. The results showed that GPX4 expression levels were significantly increased in the NE (1000 μM) + CoQ10 (1 μM) group compared to the NE group (Fig. 3E, F). Overall, these findings suggest that CoQ10 partially rescues lipid peroxidation and ferroptosis induced by NE.

**Figure 3.**
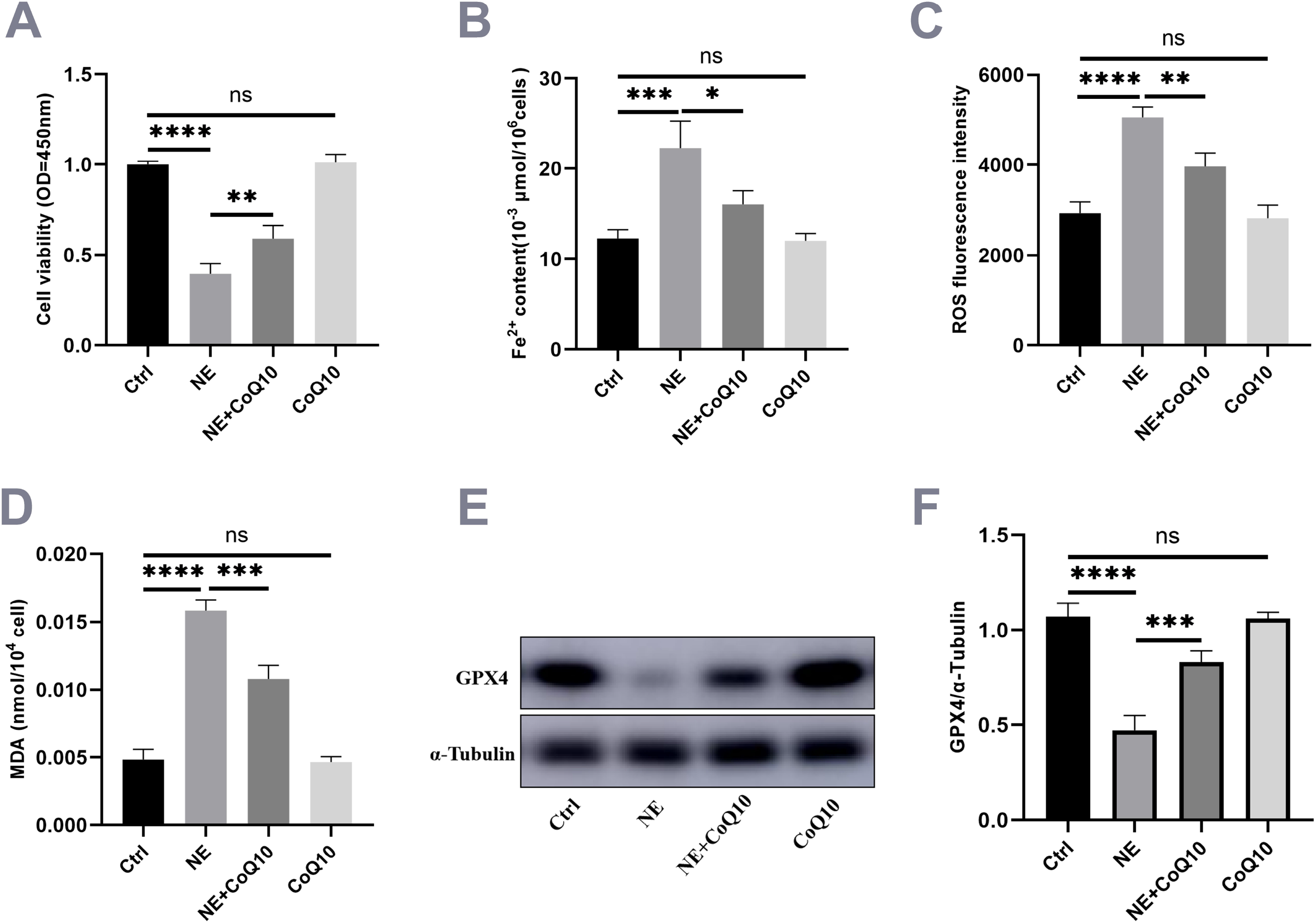
The decreased cell viability and the occurrence of ferroptosis induced by NE can be rescued by CoQ10 in KGN cells. **A**. CCK8 assay showed that KGN cell viability decreased after treatment with NE (1000 μM) for 48h, while NE (1000 μM) + CoQ10 (1 μM) could rescue KGN cell viability (n = 3). **B**. Fe^2+^ assay results showed that KGN cells treated with NE (1000 μM) for 48h had an increased Fe^2+^ content, while KGN cells treated with NE (1000 μM) + CoQ10 (1 μM) reduced Fe^2+^ content (n = 3). **C**. ROS assay showed that the ROS content increased after NE (1000μM) treatment of KGN cells and decreased after CoQ10 (1μM) supplementation for 48h (n = 3). **D**.MDA assay showed that MDA content in KGN cells increased after being treated with NE (1000μM) for 48h, while NE (1000 μM) + CoQ10 (1 μM) treated KGN cells reduced MDA levels (n = 3). **E.F**. Western blot showed that KGN cells treated with NE (1000 μM) for 48h had decreased GPX4 levels, while KGN cells treated with NE (1000 μM) + CoQ10(1 μM) increased GPX4 levels (n = 3).The data were presented as mean ± S.E.M (*p□<□0.05,**p < 0.01,***p < 0.001,****p < 0.0001).Ctrl, control; NE, norepinephrine; CoQ10, Coenzyme Q10; MDA, malondialdehyde; GPX4, glutathione peroxidase 4.

### 4. CoQ10 rescues the lipid peroxidation induced by NE in vivo

To investigate the potential protective effects of CoQ10 on ovarian function, mice were treated separately with saline, NE, CoQ10, and a combination of NE and CoQ10 (Fig. 4A). The Ferrous ion (Fe^2+^) assay results showed that the content of Fe^2+^ in the NE + CoQ10 group was significantly lower than that in the NE group (Fig. 4B). Similarly, the ROS assay results indicated that the content of Fe^2+^ in the NE + CoQ10 group was significantly lower than that in the NE group (Fig. 4C). Additionally, the examination of MDA content revealed a significant reduction in the NE + CoQ10 group compared to the NE group (Fig. 4D). Furthermore, the expression levels of GPX4 in the ovaries of the NE + CoQ10 group were increased compared to the NE group (Fig. 4E, F). Transcriptome analysis showed differential expression of genes related to ferroptosis in the ovaries of NE-treated mice compared to control mice. Importantly, the NE+CoQ10 group exhibited a rescue of the differential expression of these genes related to ferroptosis caused by NE (Fig. 4G). These findings suggest that CoQ10 can, to a certain extent, restore the lipid peroxidation of ovarian tissues caused by NE.

**Figure 4.**
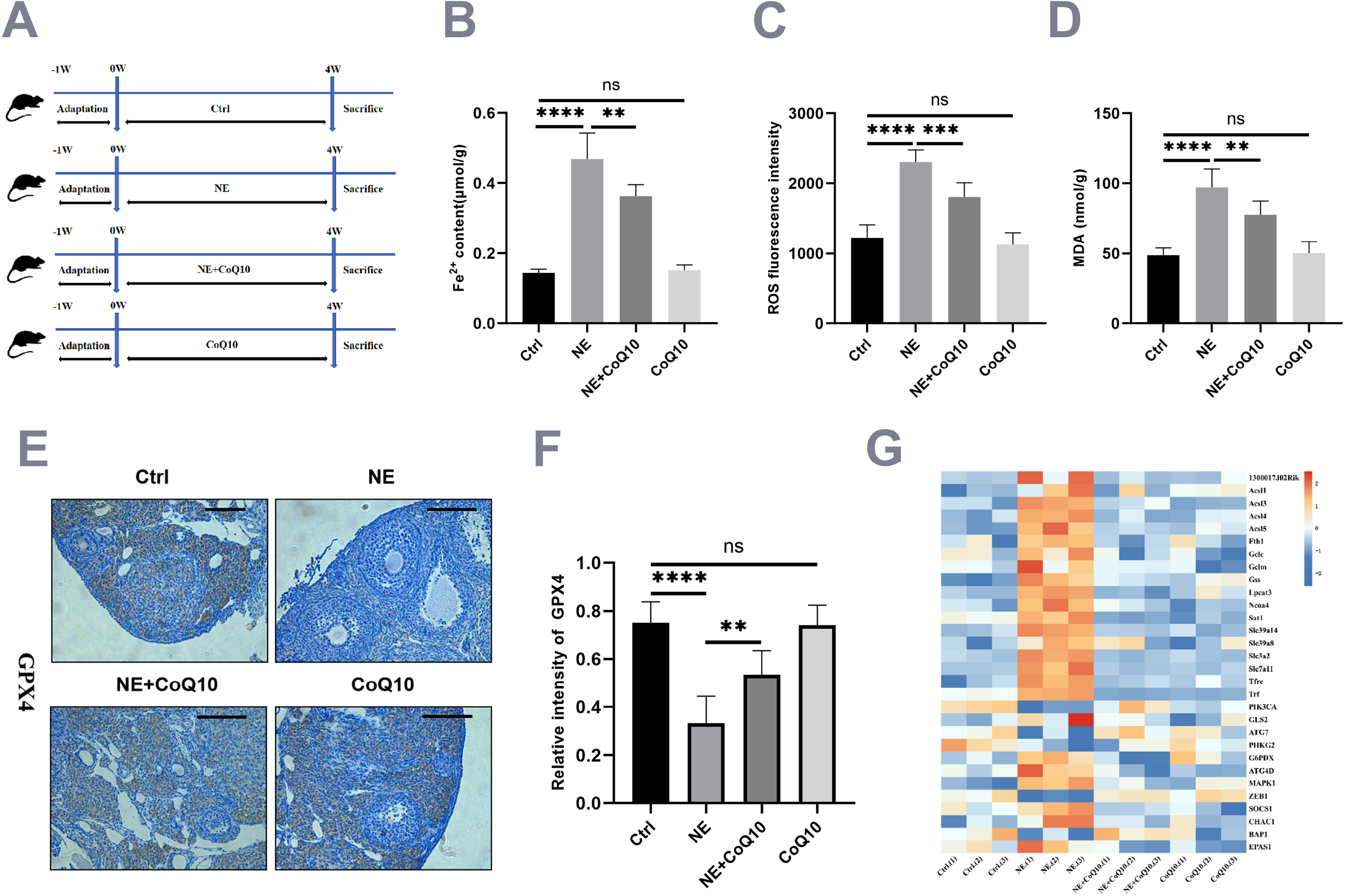
CoQ10 can rescue ovarian ferroptosis induced by NE. **A**. Experimental design of the mouse model. Female C57BL/6 mice at 8 weeks of age were randomly divided into four groups. Ctrl (saline, Ip) group, NE (1mg/kg/day, Ip) group, NE (1mg/kg/d, Ip) + CoQ10 (100mg/kg/day, feeding) group, CoQ10 (100mg/kg/day, feeding) group. The mice were treated for 4 weeks(n=3). **B**. Fe^2+^ assay results showed that Fe^2+^ was decreased in the NE+CoQ10 group compared with the NE group (n = 6). **C**. ROS assay results showed that ROS content was decreased in the NE + CoQ10 group compared with the NE group (n = 6). **D. MDA** assay showed that MDA was decreased in the NE + CoQ10 group compared with the NE group (n = 6). **E.F**.IHC results showed that GPX4 was increased in the NE + CoQ1O group compared with the NE group (n = 6, bar=100 μm). **G**. RNA-sequencing results showed that compared with the NE group, the NE + CoQ10 group could rescue the differential expression of genes related to ferroptosis caused by NE (n = 3). The data were presented as mean ± S.E.M (*p□<□0.05, **p < 0.01,***p < 0.001,****p < 0.0001).Ctrl, control; NE, norepinephrine; CoQ10, Coenzyme Q10; MDA, malondialdehyde; GPX4, glutathione peroxidase 4;Ip, intraperitoneal injection.

### 5. Ovarian dysfunction induced by NE can be rescued by CoQ10 in vivo

To further investigate the protective effect of CoQ10 on ovarian function, we examined ovarian morphology, serum hormone levels, estrous cycle, and follicle count in different groups, including the control group, NE group, NE + CoQ10 group, and CoQ10 group. We observed that the weight of ovaries in the NE + CoQ10 group increased compared to the NE group (Fig. 5E). The estrus cycle of mice was assessed by vaginal smear, revealing that NE treatment disrupted the mouse estrus cycle, while CoQ10 restored the disturbance induced by NE (Fig. 5A, B). The results of follicle count showed that the number of primordial follicles, primary follicles, secondary follicles, and antral follicles increased, whereas the number of atretic follicles decreased in the NE + CoQ10 group compared to the NE group (Fig. 5C, D). Furthermore, the levels of AMH and E2 in the NE + CoQ10 group increased compared to the NE group (Fig. 5F, G), while FSH levels decreased in the NE + CoQ10 group compared to the NE group (Fig. 5H). Overall, these findings demonstrate that CoQ10 significantly improved ovarian dysfunction induced by NE(Fig. 6).

**Figure 5.**
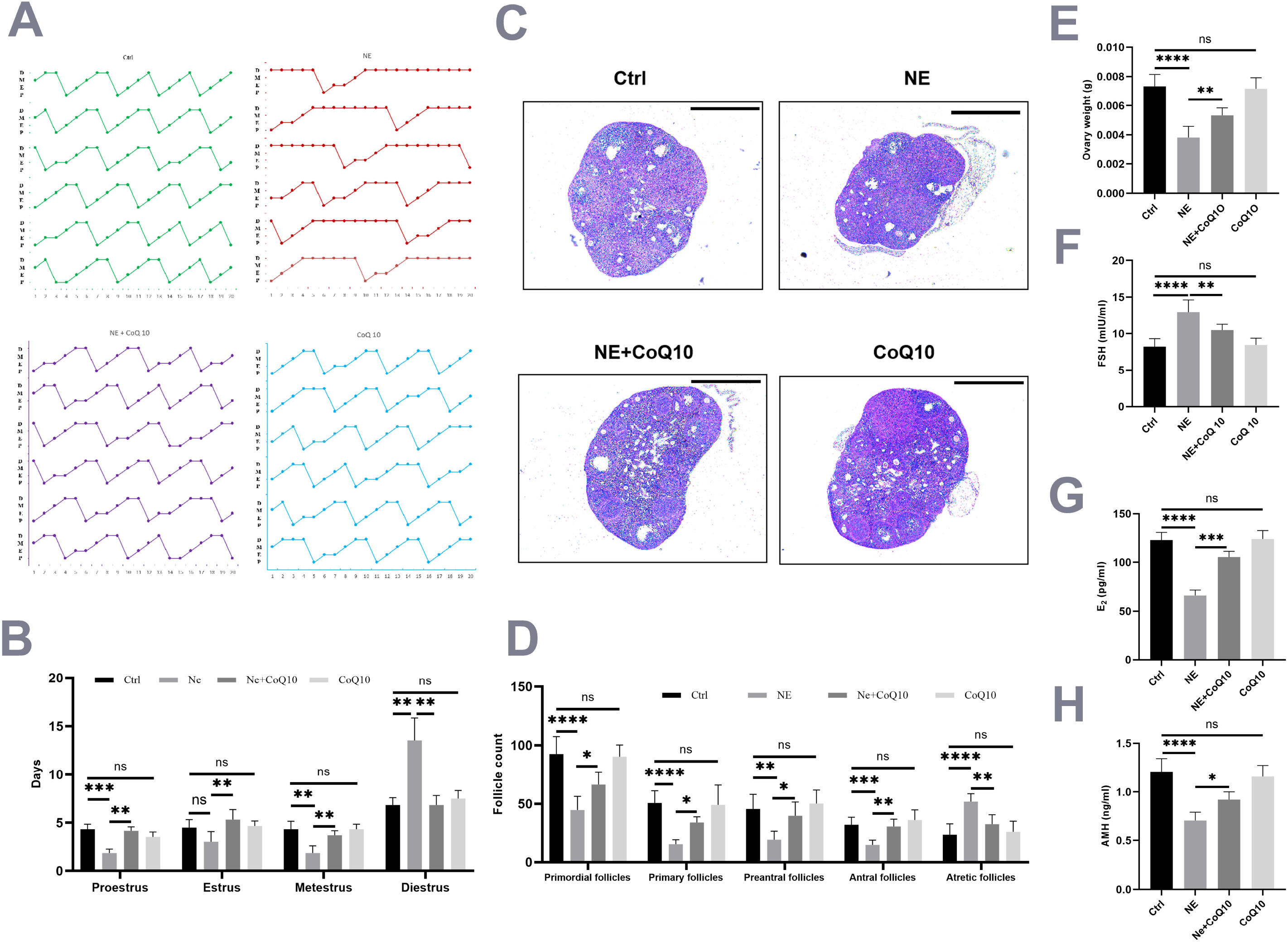
CoQ10 rescues ovarian dysfunction caused by NE. **A.B**. Vaginal smear test was used to detect the estrous cycle, and the results showed that the estrus cycle was improved in the NE + CoQ10 group compared with the NE group (n = 6). **C.D**. Compared with the NE group, the number of primordial follicles, primary follicles, secondary follicles, and antral follicles increased, and the number of atretic follicles were decreased in the NE + CoQ10 group by follicle counting (n = 6, bar=500 μm). **E**. Ovarian weight was increased in the NE + CoQ10 group compared with the NE group. (n = 6). **F.G.H**. ELISA results showed that AMH and E2 were increased and FSH was decreased in the NE + CoQ10 group compared with the NE group (n = 6). Data were presented as means ± S.E.M (*p□<□0.05,**p < 0.01,***p < 0.001,****p < 0.0001).Ctrl, control; NE, norepinephrine; CoQ10, Coenzyme Q10; ELISA, Enzyme-linked immunosorbent assay.

**Figure 6.**
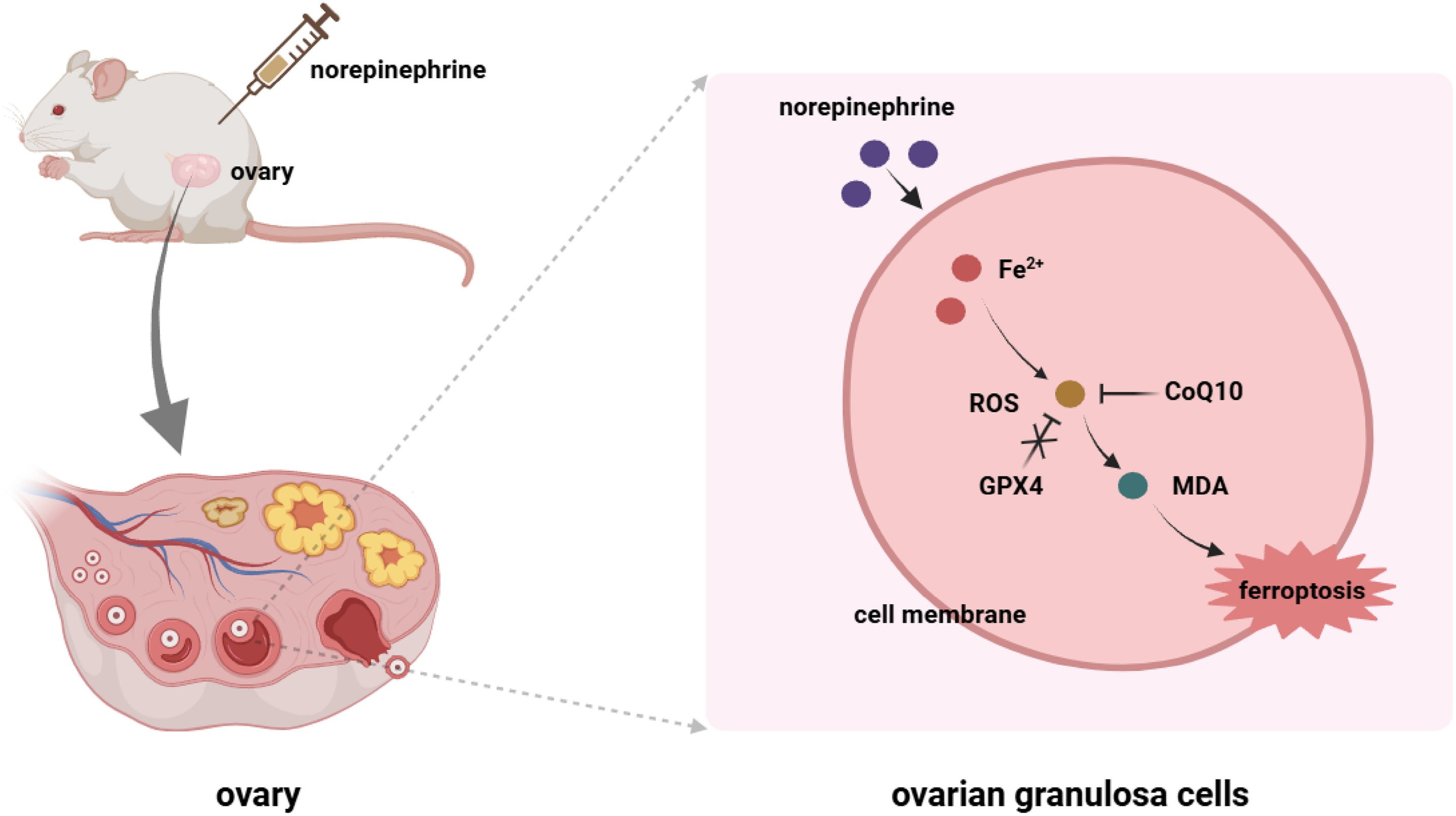
Mechanism of norepinephrine (NE) inducing ferroptosis leading to ovarian dysfunction. Excessive NE causes an increase in Fe2+, ROS, and malondialdehyde (MDA) content in ovarian granulosa cells, accompanied by a decrease in glutathione peroxidase 4 (GPX4) content, leading to ferroptosis in ovarian tissue, which can be rescued by Coenzyme Q10(CoQ10).

## Discussion

In this study, we established an animal model by intraperitoneally injecting NE into mice. Compared to the control group, the NE-treated mice exhibited a disrupted estrous cycle, characterized by prolonged diestrus and shortened estrus. Additionally, the NE group showed significantly decreased levels of AMH and E2, along with a significant increase in FSH levels. The ovaries of NE-treated mice had significantly lower weights and fewer primordial, primary, secondary, and antral follicles, but an increased number of atretic follicles. RNA sequencing results further revealed a significant enrichment of ferroptosis-related genes in the NE-treated ovaries. In vivo and in vitro experiments confirmed significant iron accumulation and ferroptosis, suggesting that NE induces ovarian dysfunction in mice by promoting ferroptosis in the ovaries. This study is the first to report that NE induces ferroptosis in the ovary, which may contribute to ovarian dysfunction. CoQ10, as an antioxidant, can protect ovarian function by inhibiting ferroptosis.

NE is a neurotransmitter released during stress reactions, and its role has been increasingly recognized among stress hormones. Studies have shown a close relationship between NE and abnormal iron metabolism[29, 30]. In a serologic analysis of women with gestational diabetes mellitus, norepinephrine, epinephrine, and transferrin were found to be positively correlated[30]. NE promotes the generation of ROS and converts cis-aconitase into iron regulatory protein-1 in the cytoplasm. It also binds to iron-responsive elements in the transferrin receptor and the non-coding region of ferritin to increase iron content in mitochondria[31]. NE has been shown to promote the growth of actinobacillus pleuropneumoniae by regulating iron uptake and metabolism[32]. Furthermore, NE can chelate iron and efficiently transport it into pseudomonas aeruginosa cells via tonb-dependent transporters, promoting bacterial growth. NE can also affect the function of transferrin, potentially leading to nervous system dysfunction[33]. The adhesion of streptococcus pneumoniae to lung epithelial cells is iron-dependent, and NE has been shown to reduce the invasiveness of streptococcus pneumoniae by binding iron[34]. Additionally, NE increases the expression of hepcidin in the liver of mice through the adrenergic receptor α1 and STAT3 pathways. The increase in hepcidin in the liver reduces the level of ferroportin 1 in the spleen, potentially leading to increased iron accumulation in the spleen [29]. These studies suggest that NE can disrupt iron metabolism and create conditions conducive to ferroptosis. In our study, we observed a significant increase in iron accumulation in mouse ovaries after treatment with NE. Similarly, the iron content in KGN cells also significantly increased after NE treatment. These results indicate that NE can induce disturbances in iron metabolism in the ovary and promote iron deposition.

Researchers have become increasingly interested in the relationship between ferroptosis and ovarian function. Ferroptosis is a form of cell death driven by iron-dependent phospholipid peroxidation, which plays a crucial role in the aging process of organisms[35]. Dysfunctional iron metabolism leads to iron accumulation and subsequent ferroptosis, resulting in cell death[36]. MiR-93-5p has been shown to promote apoptosis and ferroptosis in polycystic ovary syndrome granulosa cells through the NF-kB signaling pathway[22]. NEDD4L promotes ferroptosis in ovarian granulosa cells by facilitating GPX4 ubiquitination and degradation, contributing to the development of PCOS[37]. n-3 PUFA inhibits the proliferation of ovarian granulosa cells and promotes ferroptosis of granulosa cells in PCOS mice through YAP1-NRF2 interaction[38]. Cisplatin induces iron accumulation in ovarian tissue and increases the expression of SP1, which can bind to the promoter of ACSL4 and increase ACSL4 transcription to induce ferroptosis in granulosa cells[39]. Transferrin receptor-mediated reactive oxygen species have also been shown to promote ferroptosis in KGN cells by regulating the NADPH oxidase 1/PTEN-induced kinase 1 long-chain family member 4 signaling pathway[23]. In our study, we found that NE can promote the accumulation of iron in the ovary, which in turn drives the oxidation of unsaturated fatty acids by ROS. Additionally, the expression of GPX4, a key gene against ferroptosis, was decreased, indicating that the downregulation of GPX4 might make ovaries more susceptible to ferroptosis.

Inducers and inhibitors of ferroptosis play important roles in ovarian function. By regulating the activity of TET Ferrostatin - 1 reduces homocysteine and DNA methylation ovarian granulosa cell damage induced by homocysteine[40]. Metformin inhibits ferroptosis by regulating SIRT3/AMPK/mTOR pathway, thus improving body weight and metabolic disorders in PCOS mice, and also improving ovarian dysfunction in PCOS mice[41]. Cisplatin induces ovarian granulosa cells to produce excessive superoxide to induce mitochondrial dysfunction and trigger lipid peroxidation leading to ferroptosis. N-acetylcysteine can inhibit ferroptosis by reducing intracellular ROS levels and enhancing antioxidant capacity to improve ovarian function[42]. Spermidine has been found to attenuate ovarian injury by inhibiting oxidative stress and ferroptosis through the Nrf2/HO-1/GPX4 pathway and Akt/FHC/ACSL4 pathway[43]. Pterostilbene has been shown to ameliorate oxidative damage and ferroptosis in human ovarian granulosa cells by regulating the Nrf2/HO-1 pathway [44]. Hyperandrogenism can induce decreased viability of KGN cells, inhibited GPX4 and SLC7A11 expression, increased ACSL4 expression, increased MDA level, ROS accumulation, and increased lipid peroxidation, thereby promoting ferroptosis of KGN cells. Calcitriol can rescue hyperandrogenism induced ferroptosis of KGN cells[45].

CoQ10 is a vital compound that naturally occurs in almost every cell of the human body. It plays a crucial role in electron transport within the mitochondrial membrane during the cell’s aerobic respiration process. Adequate levels of CoQ10 are necessary for cellular respiration and ATP production. Moreover, CoQ10 is present in both HDL and LDL, where it serves an antioxidant function[46]. Studies have shown that CoQ10 can reduce ROS levels and the number of atretic follicles in mice treated with cyclophosphamide through its antioxidant effects[47]. It has also been found to improve fertility in female mice deficient in superoxide dismutase 1[48]. Clinical studies have demonstrated that CoQ10 can enhance ovarian function in patients with polycystic ovary syndrome (PCOS) by reducing insulin resistance, decreasing testosterone levels, and improving blood lipid profiles[49]. Additionally, CoQ10 supplementation has shown beneficial effects on inflammation and endothelial dysfunction in overweight and obese PCOS patients[50]. CoQ10 has also been found to improve markers of oxidative stress damage in the plasma of PCOS patients[51]. In a randomized controlled trial, CoQ10 supplementation in infertile patients increased clinical pregnancy rates by improving mitochondrial function in oocytes during in vitro maturation [52].

GPX4 is a critical endogenous antioxidant enzyme that catalyzes the reaction between reduced glutathione (GSH) and lipid hydroperoxides on the cell membrane, preventing the formation of detrimental lipid free radicals and maintaining cell membrane integrity. Decreased expression of GPX4 leads to ineffective removal of excess lipid peroxides, triggering a ferroptosis pathway dependent on ferrous ions, ultimately resulting in cell death[53]. Cells have evolved at least two defense mechanisms to inhibit ferroptosis: one is through GPX4, which uses reduced glutathione to detoxify lipid peroxides and inhibit ferroptosis. Another mechanism involves ferroptosis suppressor protein 1, functioning as an oxidoreductase that reduces ubiquinone (CoQ) to ubiquinol (CoQH2) primarily at the plasma membrane. CoQH2 then acts as a lipophilic radical-trapping antioxidant to detoxify lipid peroxyl radicals. CoQ10, with its lipophilic free radical-trapping activity on the plasma membrane, can act as an endogenous inhibitor of ferroptosis[54]. Therefore, CoQ10 likely plays a crucial role in inhibiting ferroptosis. In our study, we found that CoQ10 played a role in inhibiting ferroptosis induced by NE. We observed increased ferrous ion content, ROS content, and MDA content, along with decreased GPX4 levels in KGN cells and mouse ovaries treated with NE. However, CoQ10 treatment reduced ferrous ion, ROS, and MDA content, while increasing GPX4 levels in both cell lines and mouse ovaries exposed to NE. These findings suggest that CoQ10 plays a significant role in inhibiting NE-induced ovarian ferroptosis.

However, this study has some limitations. Firstly, the molecular mechanism of NE-induced ferroptosis in ovarian tissues needs further elucidation. Secondly, the mechanisms underlying the protective effect of CoQ10 on ovarian dysfunction induced by NE require further exploration. Further studies will provide robust evidence supporting CoQ10 as a strategy to rescue ovarian dysfunction. Thirdly, the effects of ferroptosis inhibitors on NE-induced ferroptosis and ovarian dysfunction need to be further explored. In conclusion, our study demonstrated that NE can induce ferroptosis in the mouse ovary, ultimately leading to ovarian dysfunction. CoQ10 could serve as a strategy to rescue ovarian dysfunction induced by chronic stress.

## Author contribution statement

DML and HQH conceived the project. LCW and CQX assisted with sample collection and performed the experiments. HQH performed the experiments, analysis of the data, and wrote the manuscript. DML, QW, and HQH contributed to the revised manuscript. All authors read through the manuscript and approved the final version.

## References

[1] B. Ishizuka, Current Understanding of the Etiology, Symptomatology, and Treatment Options in Premature Ovarian Insufficiency (POI), Frontiers in endocrinology 12 (2021) 626924.

[2] C.A. Stuenkel, A. Gompel, Primary Ovarian Insufficiency, The New England journal of medicine 388(2) (2023) 154–163.

[3] M. Li, Y. Zhu, J. Wei, L. Chen, S. Chen, D. Lai, The global prevalence of premature ovarian insufficiency: a systematic review and meta-analysis, Climacteric 26(2) (2023) 95–102.

[4] S. Palomba, J. Daolio, S. Romeo, F.A. Battaglia, R. Marci, G.B. La Sala, Lifestyle and fertility: the influence of stress and quality of life on female fertility, Reproductive biology and endocrinology : RB&E 16(1) (2018) 113.

[5] K.L. Rooney, A.D. Domar, The relationship between stress and infertility, Dialogues in clinical neuroscience 20(1) (2018) 41–47.

[6] L.A. Pasch, S.R. Holley, M.E. Bleil, D. Shehab, P.P. Katz, N.E. Adler, Addressing the needs of fertility treatment patients and their partners: are they informed of and do they receive mental health services?, Fertility and sterility 106(1) (2016) 209–215.e2.

[7] J.G. Schenker, D. Meirow, E. Schenker, Stress and human reproduction, European journal of obstetrics, gynecology, and reproductive biology 45(1) (1992) 1–8.

[8] J. Sun, Y. Fan, Y. Guo, H. Pan, C. Zhang, G. Mao, Y. Huang, B. Li, T. Gu, L. Wang, Q. Zhang, Q. Wang, Q. Zhou, B. Li, D. Lai, Chronic and Cumulative Adverse Life Events in Women with Primary Ovarian Insufficiency: An Exploratory Qualitative Study, Frontiers in endocrinology 13 (2022) 856044.

[9] Y.M. Ulrich-Lai, J.P. Herman, Neural regulation of endocrine and autonomic stress responses, Nature reviews. Neuroscience 10(6) (2009) 397–409.

[10] X.Y. Fu, H.H. Chen, N. Zhang, M.X. Ding, Y.E. Qiu, X.M. Pan, Y.S. Fang, Y.P. Lin, Q. Zheng, W.Q. Wang, Effects of chronic unpredictable mild stress on ovarian reserve in female rats: Feasibility analysis of a rat model of premature ovarian failure, Molecular medicine reports 18(1) (2018) 532–540.

[11] A. Keremu, A. Yaoliwasi, M. Tuerhong, N. Kadeer, Heyi, A. Yiming, X. Yilike, Research on the establishment of chronic stress-induced premature ovarian failure the rat model and effects of Chinese medicine Muniziqi treatment, Molecular reproduction and development 86(2) (2019) 175–186.

[12] R.R. Chen, J. Wang, M. Zhang, Q.Q. Kong, G.Y. Sun, C.H. Jin, M.J. Luo, J.H. Tan, Restraint stress of female mice during oocyte development facilitates oocyte postovulatory aging, Aging 14(22) (2022) 9186–9199.

[13] G. Russell, S. Lightman, The human stress response, Nat Rev Endocrinol 15(9) (2019) 525–534.

[14] S. Saller, J. Merz-Lange, S. Raffael, S. Hecht, R. Pavlik, C. Thaler, D. Berg, U. Berg, L. Kunz, A. Mayerhofer, Norepinephrine, active norepinephrine transporter, and norepinephrine-metabolism are involved in the generation of reactive oxygen species in human ovarian granulosa cells, Endocrinology 153(3) (2012) 1472–83.

[15] N. Musalı, B. Özmen, Y.E. Şükür, B. Ergüder, C.S. Atabekoğlu, M. Sönmezer, B. Berker, R. Aytaç, Follicular fluid norepinephrine and dopamine concentrations are higher in polycystic ovary syndrome, Gynecological endocrinology : the official journal of the International Society of Gynecological Endocrinology 32(6) (2016) 460–3.

[16] L. Wang, C. Zhou, J. Sun, Q. Zhang, D. Lai, Glutamine and norepinephrine in follicular fluid synergistically enhance the antioxidant capacity of human granulosa cells and the outcome of IVF-ET, Sci Rep 12(1) (2022) 9936.

[17] S.J. Dixon, K.M. Lemberg, M.R. Lamprecht, R. Skouta, E.M. Zaitsev, C.E. Gleason, D.N. Patel, A.J. Bauer, A.M. Cantley, W.S. Yang, B. Morrison, 3rd, B.R. Stockwell, Ferroptosis: an iron-dependent form of nonapoptotic cell death, Cell 149(5) (2012) 1060–72.

[18] J. Li, F. Cao, H.L. Yin, Z.J. Huang, Z.T. Lin, N. Mao, B. Sun, G. Wang, Ferroptosis: past, present and future, Cell death & disease 11(2) (2020) 88.

[19] X. Jiang, B.R. Stockwell, M. Conrad, Ferroptosis: mechanisms, biology and role in disease, Nature reviews. Molecular cell biology 22(4) (2021) 266–282.

[20] P.H. Lin, C.J. Li, L.T. Lin, W.P. Su, J.J. Sheu, Z.H. Wen, J.T. Cheng, K.H. Tsui, Unraveling the Clinical Relevance of Ferroptosis-Related Genes in Human Ovarian Aging, Reprod Sci 30(12) (2023) 3529–3536.

[21] F. Wang, Y. Liu, F. Ni, J. Jin, Y. Wu, Y. Huang, X. Ye, X. Shen, Y. Ying, J. Chen, R. Chen, Y. Zhang, X. Sun, S. Wang, X. Xu, C. Chen, J. Guo, D. Zhang, BNC1 deficiency-triggered ferroptosis through the NF2-YAP pathway induces primary ovarian insufficiency, Nat Commun 13(1) (2022) 5871.

[22] W. Tan, F. Dai, D. Yang, Z. Deng, R. Gu, X. Zhao, Y. Cheng, MiR-93-5p promotes granulosa cell apoptosis and ferroptosis by the NF-kB signaling pathway in polycystic ovary syndrome, Frontiers in immunology 13 (2022) 967151.

[23] L. Zhang, F. Wang, D. Li, Y. Yan, H. Wang, Transferrin receptor-mediated reactive oxygen species promotes ferroptosis of KGN cells via regulating NADPH oxidase 1/PTEN induced kinase 1/acyl-CoA synthetase long chain family member 4 signaling, Bioengineered 12(1) (2021) 4983–4994.

[24] F. Li, Y. Wang, M. Xu, N. Hu, J. Miao, Y. Zhao, L. Wang, Single-nucleus RNA Sequencing reveals the mechanism of cigarette smoke exposure on diminished ovarian reserve in mice, Ecotoxicol Environ Saf 245 (2022) 114093.

[25] X. Fang, H. Ardehali, J. Min, F. Wang, The molecular and metabolic landscape of iron and ferroptosis in cardiovascular disease, Nat Rev Cardiol 20(1) (2023) 7–23.

[26] M. Dodson, R. Castro-Portuguez, D.D. Zhang, NRF2 plays a critical role in mitigating lipid peroxidation and ferroptosis, Redox biology 23 (2019) 101107.

[27] D. Liang, Y. Feng, F. Zandkarimi, H. Wang, Z. Zhang, J. Kim, Y. Cai, W. Gu, B.R. Stockwell, X. Jiang, Ferroptosis surveillance independent of GPX4 and differentially regulated by sex hormones, Cell 186(13) (2023) 2748–2764.e22.

[28] K. Bersuker, J.M. Hendricks, Z. Li, L. Magtanong, B. Ford, P.H. Tang, M.A. Roberts, B. Tong, T.J. Maimone, R. Zoncu, M.C. Bassik, D.K. Nomura, S.J. Dixon, J.A. Olzmann, The CoQ oxidoreductase FSP1 acts parallel to GPX4 to inhibit ferroptosis, Nature 575(7784) (2019) 688–692.

[29] W.N. Kong, Y. Cui, Y.J. Fu, Y. Lei, Y. Ci, Y. Bao, S. Zhao, L. Xie, Y.Z. Chang, S.E. Zhao, The α1-adrenergic receptor is involved in hepcidin upregulation induced by adrenaline and norepinephrine via the STAT3 pathway, Journal of cellular biochemistry 119(7) (2018) 5517–5527.

[30] Y. Feng, Q. Feng, Y. Lv, X. Song, H. Qu, Y. Chen, The relationship between iron metabolism, stress hormones, and insulin resistance in gestational diabetes mellitus, Nutrition & diabetes 10(1) (2020) 17.

[31] N. Tapryal, G.V. Vivek, C.K. Mukhopadhyay, Catecholamine stress hormones regulate cellular iron homeostasis by a posttranscriptional mechanism mediated by iron regulatory protein: implication in energy homeostasis, The Journal of biological chemistry 290(12) (2015) 7634–46.

[32] L. Li, Z. Chen, W. Bei, Z. Su, Q. Huang, L. Zhang, H. Chen, R. Zhou, Catecholamines promote Actinobacillus pleuropneumoniae growth by regulating iron metabolism, PloS one 10(4) (2015) e0121887.

[33] Q. Perraud, L. Kuhn, S. Fritsch, G. Graulier, V. Gasser, V. Normant, P. Hammann, I.J. Schalk, Opportunistic use of catecholamine neurotransmitters as siderophores to access iron by Pseudomonas aeruginosa, Environmental microbiology 24(2) (2022) 878–893.

[34] X.F. Gonzales, G. Castillo-Rojas, A.I. Castillo-Rodal, E. Tuomanen, Y. López-Vidal, Catecholamine norepinephrine diminishes lung epithelial cell adhesion of Streptococcus pneumoniae by binding iron, Microbiology (Reading, England) 159(Pt 11) (2013) 2333–2341.

[35] M. Mazhar, A.U. Din, H. Ali, G. Yang, W. Ren, L. Wang, X. Fan, S. Yang, Implication of ferroptosis in aging, Cell death discovery 7(1) (2021) 149.

[36] W.S. Yang, B.R. Stockwell, Ferroptosis: Death by Lipid Peroxidation, Trends Cell Biol 26(3) (2016) 165–176.

[37] H. Tang, X. Jiang, Y. Hua, H. Li, C. Zhu, X. Hao, M. Yi, L. Li, NEDD4L facilitates granulosa cell ferroptosis by promoting GPX4 ubiquitination and degradation, Endocrine connections 12(4) (2023).

[38] P. Zhang, Y. Pan, S. Wu, Y. He, J. Wang, L. Chen, S. Zhang, H. Zhang, Y. Zhao, L. Niu, M. Gan, Y. Wang, L. Shen, L. Zhu, n-3 PUFA Promotes Ferroptosis in PCOS GCs by Inhibiting YAP1 through Activation of the Hippo Pathway, Nutrients 15(8) (2023).

[39] S. Wang, X. Li, J. Li, A. Wang, F. Li, H. Hu, T. Long, X. Pei, H. Li, F. Zhong, F. Zhu, Inhibition of cisplatin-induced Acsl4-mediated ferroptosis alleviated ovarian injury, Chem Biol Interact 387 (2024) 110825.

[40] Q. Shi, R. Liu, L. Chen, Ferroptosis inhibitor ferrostatinlZI1 alleviates homocysteinelZIinduced ovarian granulosa cell injury by regulating TET activity and DNA methylation, Molecular medicine reports 25(4) (2022).

[41] Q. Peng, X. Chen, X. Liang, J. Ouyang, Q. Wang, S. Ren, H. Xie, C. Wang, Y. Sun, X. Wu, H. Liu, C. Hei, M. Sun, Q. Chang, X. Liu, G. Li, R. He, Metformin improves polycystic ovary syndrome in mice by inhibiting ovarian ferroptosis, Frontiers in endocrinology 14 (2023) 1070264.

[42] S. Zhang, Q. Liu, M. Chang, Y. Pan, B.H. Yahaya, Y. Liu, J. Lin, Chemotherapy impairs ovarian function through excessive ROS-induced ferroptosis, Cell death & disease 14(5) (2023) 340.

[43] C. Niu, D. Jiang, Y. Guo, Z. Wang, Q. Sun, X. Wang, W. Ling, X. An, C. Ji, S. Li, H. Zhao, B. Kang, Spermidine suppresses oxidative stress and ferroptosis by Nrf2/HO-1/GPX4 and Akt/FHC/ACSL4 pathway to alleviate ovarian damage, Life sciences 332 (2023) 122109.

[44] X. Chen, Q.L. Song, Z.H. Li, R. Ji, J.Y. Wang, M.L. Cao, X.F. Mu, Y. Zhang, D.Y. Guo, J. Yang, Pterostilbene ameliorates oxidative damage and ferroptosis in human ovarian granulosa cells by regulating the Nrf2/HO-1 pathway, Archives of biochemistry and biophysics 738 (2023) 109561.

[45] Y. Jiang, J. Yang, K. Du, K. Luo, X. Yuan, F. Hua, 1,25-Dihydroxyvitamin D3 alleviates hyperandrogen-induced ferroptosis in KGN cells, Hormones (Athens, Greece) 22(2) (2023) 273–280.

[46] J. Garrido-Maraver, M.D. Cordero, M. Oropesa-Avila, A.F. Vega, M. de la Mata, A.D. Pavon, E. Alcocer-Gomez, C.P. Calero, M.V. Paz, M. Alanis, I. de Lavera, D. Cotan, J.A. Sanchez-Alcazar, Clinical applications of coenzyme Q10, Frontiers in bioscience (Landmark edition) 19(4) (2014) 619–33.

[47] A. Delkhosh, M. Delashoub, A.A. Tehrani, A.M. Bahrami, V. Niazi, H. Shoorei, M. Banimohammad, H. Kalarestaghi, M. Shokoohi, A. Agabalazadeh, M. Mohaqiq, Upregulation of FSHR and PCNA by administration of coenzyme Q10 on cyclophosphamide-induced premature ovarian failure in a mouse model, Journal of biochemical and molecular toxicology 33(11) (2019) e22398.

[48] N. Ishii, T. Homma, J. Lee, H. Mitsuhashi, K.I. Yamada, N. Kimura, Y. Yamamoto, A.J. Fujii, Ascorbic acid and CoQ10 ameliorate the reproductive ability of superoxide dismutase 1-deficient female mice†, Biology of reproduction 102(1) (2020) 102–115.

[49] P.H. Lin, W.P. Su, C.J. Li, L.T. Lin, J.J. Sheu, Z.H. Wen, J.T. Cheng, K.H. Tsui, Investigating the Role of Ferroptosis-Related Genes in Ovarian Aging and the Potential for Nutritional Intervention, Nutrients 15(11) (2023).

[50] S. Taghizadeh, A. Izadi, S. Shirazi, M. Parizad, B. Pourghassem Gargari, The effect of coenzyme Q10 supplementation on inflammatory and endothelial dysfunction markers in overweight/obese polycystic ovary syndrome patients, Gynecological endocrinology : the official journal of the International Society of Gynecological Endocrinology 37(1) (2021) 26–30.

[51] C. Di Segni, A. Silvestrini, R. Fato, C. Bergamini, F. Guidi, S. Raimondo, E. Meucci, D. Romualdi, R. Apa, A. Lanzone, A. Mancini, Plasmatic and Intracellular Markers of Oxidative Stress in Normal Weight and Obese Patients with Polycystic Ovary Syndrome, Experimental and clinical endocrinology & diabetes : official journal, German Society of Endocrinology [and] German Diabetes Association 125(8) (2017) 506–513.

[52] M.K. Abdulhasan, Q. Li, J. Dai, H.M. Abu-Soud, E.E. Puscheck, D.A. Rappolee, CoQ10 increases mitochondrial mass and polarization, ATP and Oct4 potency levels, and bovine oocyte MII during IVM while decreasing AMPK activity and oocyte death, Journal of assisted reproduction and genetics 34(12) (2017) 1595–1607.

[53] F. Ursini, M. Maiorino, Lipid peroxidation and ferroptosis: The role of GSH and GPx4, Free radical biology & medicine 152 (2020) 175–185.

[54] C. Mao, X. Liu, Y. Zhang, G. Lei, Y. Yan, H. Lee, P. Koppula, S. Wu, L. Zhuang, B. Fang, M.V. Poyurovsky, K. Olszewski, B. Gan, DHODH-mediated ferroptosis defence is a targetable vulnerability in cancer, Nature 593(7860) (2021) 586–590.

